# Inhibition of eukaryotic translation initiation factor 1A (eIF1A) and 3B (eIF3B) diminishes the psoriatic phenotype in two mouse models and human 3D model samples

**DOI:** 10.1101/2024.03.21.586074

**Authors:** Nicole Golob-Schwarzl, Johannes Pilic, Azelma Vrebo, Natalie Bordag, Nitesh Shirsath, Christin-Therese Müller, Amin El-Heliebi, Peter Wolf

**Author notes:** Correspondence: Nicole Golob-Schwarzl Department of Dermatology and Venereology, Medical University of Graz, Auenbruggerplatz 8, 8036 Graz, Austria.

## Abstract

**Background:** Psoriasis is a systemic inflammatory disease for which new topical treatments are required. Psoriatic inflammation is associated with overexpression of eukaryotic translation initiation factors (eIFs), which critically regulate gene expression in many important cellular processes, including proliferation, apoptosis, and differentiation. However, the exact link between overexpression of eIF and psoriasis is unknown. Here, we investigated the role of eIFs, particularly eIF1A and eIF3B, and the impact of their inhibition on the pathophysiology of psoriasis.

**Methods:** We used two mouse models reflecting the pathophysiology of psoriasis: (i) BALB/c mice topically treated with the immune activator imiquimod (IMQ) and (ii) K5.TGFβ transgenic mice. eIF1A and eIF3B were inhibited by either topical or systemic application of specific small interfering RNA (siRNA). In addition, we employed commercial human 3D psoriatic skin model samples. Importantly, *in situ* mRNA detection-based padlock probes against transcript variants of eIF1A und eIF3B was performed.

**Results:** Topical and systemic inhibition of eIF1A and eIF3B inhibited inflammation in both imiquimod and TGFß mouse models as well as in a human 3D psoriasis model. Downregulation of eIF1A and eIF3B was associated with normalization of cell proliferation, restoration of the inflammatory milieu and epidermal hyperplasia of psoriasis, and normalization of levels of proinflammatory cytokines (e.g., TNFα, IL-1b, IL-17, and IL-22) and keratinocyte differentiation markers (e.g., KRT16 and FLG).

**Conclusion:** These results reveal an imbalance in translation and emphasize the crucial role of eIF1A and eIF3B in the pathophysiology of psoriasis. Targeting eIFs opens new avenues for the development of novel therapeutic treatment strategies against psoriasis.

## Introduction

Psoriasis is a systemic inflammatory disease [1] with a polygenic predisposition that mechanistically increases the life-shortening comorbidity rate [2]. The pathogenesis of psoriasis is complex and characterized by dysregulations of skin thickness, barrier function, immune cell infiltration and function, cytokine levels, epigenetic alterations and disturbances of microbiota [1]. In our recent work [3], we observed that specific eIF4E inhibition improved the psoriasis phenotype, thus, we herein investigated eukaryotic translation initiation and its inhibition in further detail.

The initiation of eukaryotic translation is a multistep process involving ribosomes, mRNAs, tRNAs and a number of proteins called eukaryotic translation initiation factors (eIFs). During the initiation of translation, active 80S complexes are formed when small (40S) and large (60S) ribosomal subunits on the mRNA come together with the start codon and a methionyl inhibitor tRNA (Met-tRNAiMet) at the peptidyl site. In eukaryotes, at least 12 initiation factors (eIFs), consisting of about 28 polypeptides, are involved in this process, whereas in prokaryotes only three factors are required [4]. In a reconstituted translation initiation system with purified yeast components and an uncapped, unstructured mRNA [5], eIF1 and eIF1A are sufficient to promote the assembly of a 43S mRNA complex consisting of the 40S subunit, the mRNA and the ternary complex. The 43S complex is thought to scan along the mRNA until it recognizes the initiation codon, which initiates the dissociation of eIF1 and the subsequent release of phosphate by the GTPase eIF2 [5, 6]. This irreversible step is promoted by the activity of eIF5, a GTPase-activating protein [5]. Finally, eIF5B promotes dissociation of the initiation factor and assembly of the subunits to form an active 80S ribosome containing the inhibitor tRNA within the P site paired with the start codon AUG. *In vivo*, another multi-subunit factor, eIF3, plays a crucial role in the formation of the 43S complex and, together with eIF4F, eIF4B and PAB, is required to load mRNAs into the preinitiation complex. eIF1A is a small protein (17kDa) that binds with high affinity to the 40S subunit [7] and is highly conserved in all eukaryotes [4, 8]. eIF3 is the largest translation initiation complex. It consists of 13 subunits in mammals (A, B, C, D, E, F, G, H, I, K, L, M), which correspond to five nuclear subunits in the yeast Saccharomyces cerevisiae (a/Tif32, b/Prt1, c/Nip1, i/Tif34, g/Tif35) [9, 10]. Increasing evidence suggests that eIF3 plays a unique role in regulating the translation initiation process [7], translation termination [11], ribosomal recycling [12, 13] and read-through of the programmed stop codon [14, 15]. Due to the essential function of eIF3 in various physiological processes, its dysregulation has been shown to be associated with various pathological conditions, in particular with the occurrence, development and prognosis of various human cancers. Recent studies have shown that upregulation of eIF3A/ B/ C/ H/ I/ M or downregulation of eIF3E and eIF3F is related to metastasis in various cancers [16-19]. However, the treatment and prognostic role of eIF1A and eIF3B3 in psoriasis has not yet been clarified.

To utilize eIF1A and eIF3B as potential therapeutic targets in psoriasis, we used the imiquimod (IMQ) and TGFβ mouse model and a 3D human psoriasis model to investigate the functional significance of eIF1A and eIF3B in psoriatic inflammation. We specifically investigated their inhibition by siRNA as a potential therapeutic approach and found that eIF1A and eIF3B drive psoriatic inflammation and hyperplasia, while pharmaceutical inhibition of these eIFs can reduce the psoriatic phenotype. This work opens new avenues for the development of novel psoriasis treatment strategies targeting eIF1A and eIF3B.

## Results

### Knockdown of eIF1A and eIF3B reduces the proliferation of HaCaT cells

To determine the effectiveness of eIF1A knockdown in HaCaT cells, western blot analysis was performed showing significantly reduced protein levels of eIF1A after one and two days of silencing (Figure 1A). On day 3, silencing of eIF1A was not as pronounced as on days 1 and 2 (Figure 1A).

**Figure 1.**
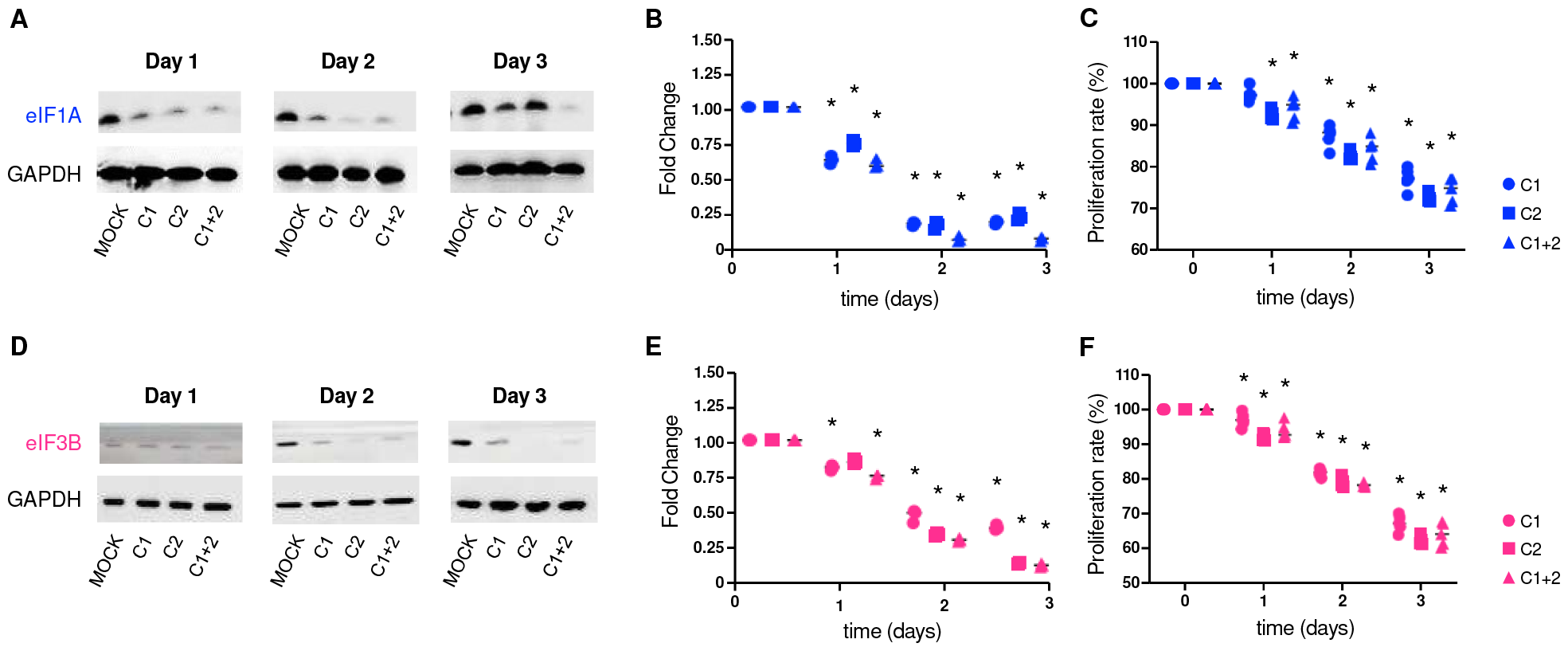
Knockdown of eIF1A and eIF3B reduces proliferation in HaCaT cells. (**A**) Immunoblot of eIF1A after siRNA transfection. (**B**) eIF1A mRNA levels after siRNA transfection. (**C**) Proliferation rate after eIF1A siRNA transfection. (**D**) Immunoblot of eIF3B after siRNA transfection. (**E**) eIF3B mRNA levels after siRNA transfection. (**F**) Proliferation rate after eIF3B siRNA transfection. ^*^ *p* < 0.05; ^**^ *p* < 0.01; ^***^ *p* < 0.001; ^****^ *p* < 0.0001.

Silencing was efficient as shown by the significant reduction in eIF1A mRNA expression observed for both eIF1A-siRNA constructs (C1 and C2) and for their combination (C1+2) by 25-30% on day 1, 75-90% on day 2 and 70-90% on day 3 (Figure 1B), compared to MOCK-treated cells. The proliferation rate was reduced by 5 - 12 % on day 1, 8 - 20 % on day 2 and 18 - 30 % on day 3 (Figure 1C).

For eIF3B silencing, western blot showed reduced bands on day 2 and day 3 and much weaker bands on day 1 (Figure 1D). eIF3B siRNA showed a significant reduction in eIF3B mRNA expression for both constructs (C-1 and C-2) and for their combination (C-1+2) by 5-30 % on day 1, 52-75 % on day 2 and 48-95 % on day 3 (Figure 1E). The proliferation rate was reduced by 2-12% on day 1, 15-25% on day 2 and 28-42% on day 3 after eIF3B silencing treatment (Figure 1F).

These results indicate that eIF1A and eIF3B are functionally relevant in the context of keratinocyte proliferation. Since psoriasis is characterized by keratinocyte proliferation, specific inhibition of eIF1A and eIF3B by siRNA could be a potential approach for the treatment of psoriasis.

### Topical and systemic application of eIF1A and eIF3B siRNA ameliorates the psoriatic phenotype in a human 3D psoriasis tissue model

A human 3D model of psoriasis was used to study the effects of topical and systemic treatment with eIF1A and eIF3B siRNA. Topical treatment with eIF1A siRNA reduced the epidermal thickness from 170 µm to 52, 92 and 68 µm after topical administration (Figure 2A) and to 103, 52 and 75 µm after systemic application in culture medium (Figure 2A) for both eIF1A siRNA constructs (C1 and C2) and for their combination (C1+2), respectively. Similarly, topical treatment with eIF3B siRNA reduced the epidermal thickness from 148 µm to 53 µm and to 51 µm after topical administration (Figure 2C), to 52 µm, 78 µm, and to 54 µm after application in culture medium (Figure 2C). RT-PCR of eIF1A and eIF3B showed significant downregulation after topical and systemic siRNA treatment for both constructs (C1 and C2) and the combination (C1+C2) (Figure 2B and Figure 2D).

**Figure 2.**
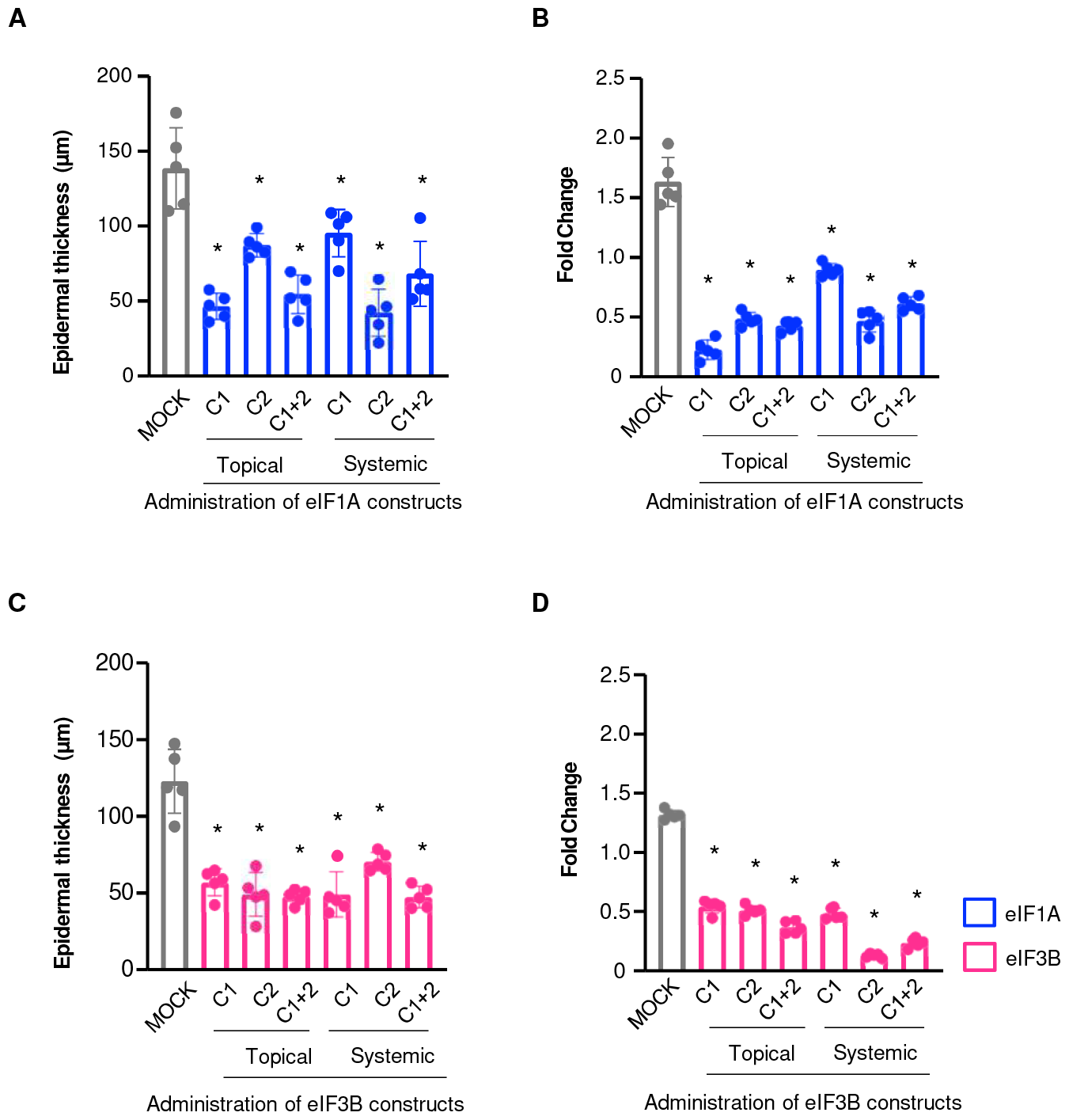
Application of eIF1A and eIF3B siRNA diminishes pathological phenotype of human 3D psoriasis tissue model. **(A)** Mean epidermal thickness after topical and systemic eIF1A treatment. **(B)** eIF1A mRNA levels after topical and systemic eIF1A siRNA transfection. **(C)** Mean epidermal thickness after topical and systemic eIF3B treatment. **(D)** eIF3B mRNA levels after topical and systemic eIF3B siRNA transfection ^*^ *p* < 0.05; ^**^ *p* < 0.01; ^***^ *p* < 0.001; ^***^ *p* < 0.0001.

Taken together, these results demonstrate that topical administration of eIF1A and eIF3B siRNA is an anti-psoriasis approach that improves both the keratinocyte compartment and inflammatory infiltrates in the epidermis.

### Topical and systemic eIF1A and eIF3B siRNA treatment reduces the macroscopic and microscopic psoriatic phenotype in IMQ and TGFß mice

To test the effects of topical and systemic eIF1A and eIF3B siRNA treatment in an *in vivo* setting, we used the well-established IMQ and TGFß psoriatic mouse models. Topical and systemic application of eIF1A and eIF3B siRNA significantly improved the psoriatic phenotype in both IMQ (Figure 3A) and TGFß (Figure 5A) mouse models. Treatment with IMQ resulted in an increase in double skinfold thickness during application from 0.55 mm on day 1 to 1.25 mm on day 7 (Figure 3B). In comparison, concomitant treatment with topical or systemic eIF1A or eIF3B siRNA resulted in a large reduction in skinfold thickness from 1.25 mm on day 1 to 0.73 mm, 0.70 mm, 0.68 mm and 0.62 mm on day 7. Treatment not only eliminated the psoriatic manifestation at the macroscopic level (Figure 3C and 5C, upper panels), but also reduced epidermal hyperproliferation and inflammatory infiltration of the skin (Figure 3C and 5C, lower two panels) in both mouse models. RT-PCR of TNFα, IL -1b, IL -17, IL -22, KRT16, S100A8 and FLG showed down-regulation after topical and systemic siRNA for both constructs in both models (Figure 3D and Figure 5D). Application of IMQ increased the average thickness of the epidermis from 50 to 98 µm, and topical and systemic treatment with eIF1A and eIF3B siRNA normalized this thickness to 52 and 58 µm, respectively (Figure 3E).

**Figure 3.**
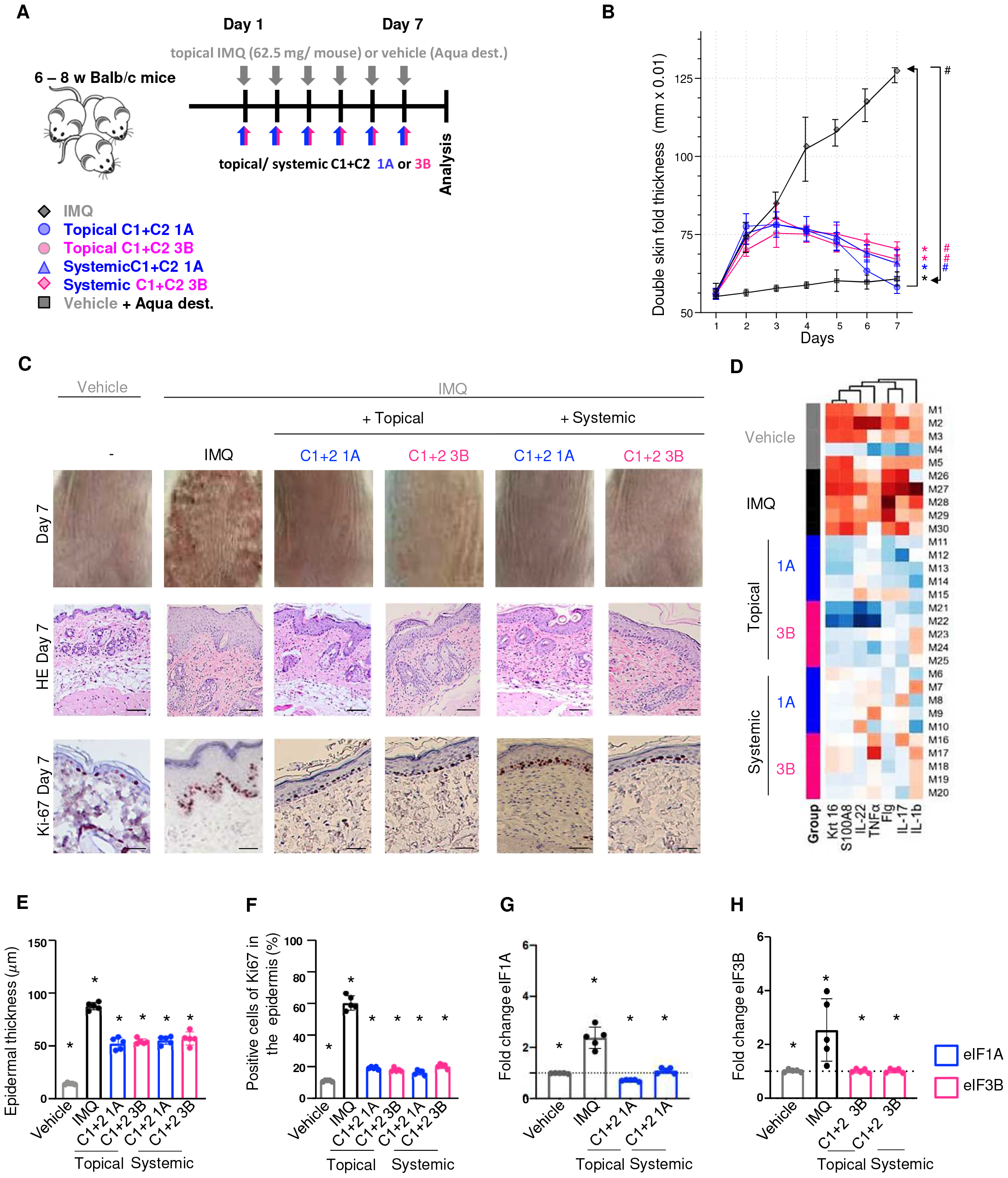
Topical and systemic administration of eIF1A and eIF3B siRNA ameliorates the psoriatic phenotype of IMQ mice. **(A)** Topical and systemic administration of specific eIF1A and eIF3B siRNA 6 h after daily IMQ treatment for seven days. **(B)** Double skinfold thickness over the time of topical and systemic eIF1A and eIF3B siRNA application. **(C)** H & E and Ki-67 staining after topical or systemic eIF1A and eIF3B siRNA treatment on day 7. **(D)** Heat map of expression of proinflammatory cytokines and keratinocyte markers (ΔΔCT from RT-PCR, z-scaled per cytokine, normalized to reference group IMQ) shown with dendrograms sorted by average distance from dendrograms of hierarchical cluster analysis. **(E)** Epidermal thickness. (**F**) Ki-67 positive stained cells in the epidermis. **(G)** mRNA expression of *eIF1A*. **(H)** mRNA expression of *eIF3B. N* = 5 per group. ^*^ *p* < 0.05; ^*^ *p* < 0.01; ^***^ *p* < 0.001; ^****^ *p* < 0.0001.

In the TGFß model, both topical and systemic eIF1A and eIF3B siRNA treatment reduced the double skinfold thickness from 1.25 mm on day 1 to 0.58 mm on day 7 (Figure 5B). Treatment with eIF1A and eIF3B siRNA reduced the average thickness of the epidermis from 104 to 52 µm (Figure 5E).

In the IMQ model, eIF1A and eIF3B siRNA reduced Ki-67 positivity topically in the epidermis from 70% to 21% and 20%, respectively, and systemically from 98% to 18% and 24%, respectively (Figure 3F). In the epidermis, eIF1A and eIF3B siRNA reduced Ki-67 positivity in the TGFß model topically from 90% to 21 and 23%, respectively, and systemically from 90% to 28 and 21%, respectively (Figure 5F).

RT-PCR showed a significant down-regulation of topical eIF1A expression below normal levels in both mouse models (Figures 3G and 5G). The mRNA expression of topical and systemic eIF1A as well as systemic eIF3B was significantly downregulated to normal levels in both mouse models (Figures 3G and 3H as well as Figures 5G and 5H).

Taken together, these results suggest that topical and systemic inhibition of eIF1A and eIF3B is effective in reducing in ameliorating psoriatic phenotypes in the IMQ and TGFß models.

### Interaction potential between eIF1A and eIF3B after topical and systemic treatment in psoriasis

It is not clear whether keratinocytes possess the transcription machinery to produce eIFs. Furthermore, it is not entirely clear how the interaction distribution of eIF1A and eIF3B behaves in keratinocytes. Therefore, we used a highly specific approach *in situ* sequencing approach for mRNA detection based on padlock probes targeting transcript variants of eIF1A and eIF3B (Figure 4 and Figure 6) (PubMed ID: 32990747). We observed positive signals for eIF1A and eIF3B in epidermal keratinocytes of IMQ and TGFß mouse models (Figure 4 and Figure 6). The expression of eIF1A and eIF3B was significantly higher in the IMQ and TGFß TG group than in the vehicle and TGFß WT group. After topical and systemic application of eIF1A siRNA in IMQ and TGFß TG mice, the positive signals of eIF1A were lower compared to eIF3B in keratinocytes of the epidermis (Figure 4 and Figure 6). After topical ans systemic application of eIF3B siRNA, we observed less positive signaling of eIF3B in IMQ and TGFß TG mice (Figure 4 and Figure 6). These results suggest that keratinocytes possess the transcription machinery to produces eIFs.

**Figure 4.**
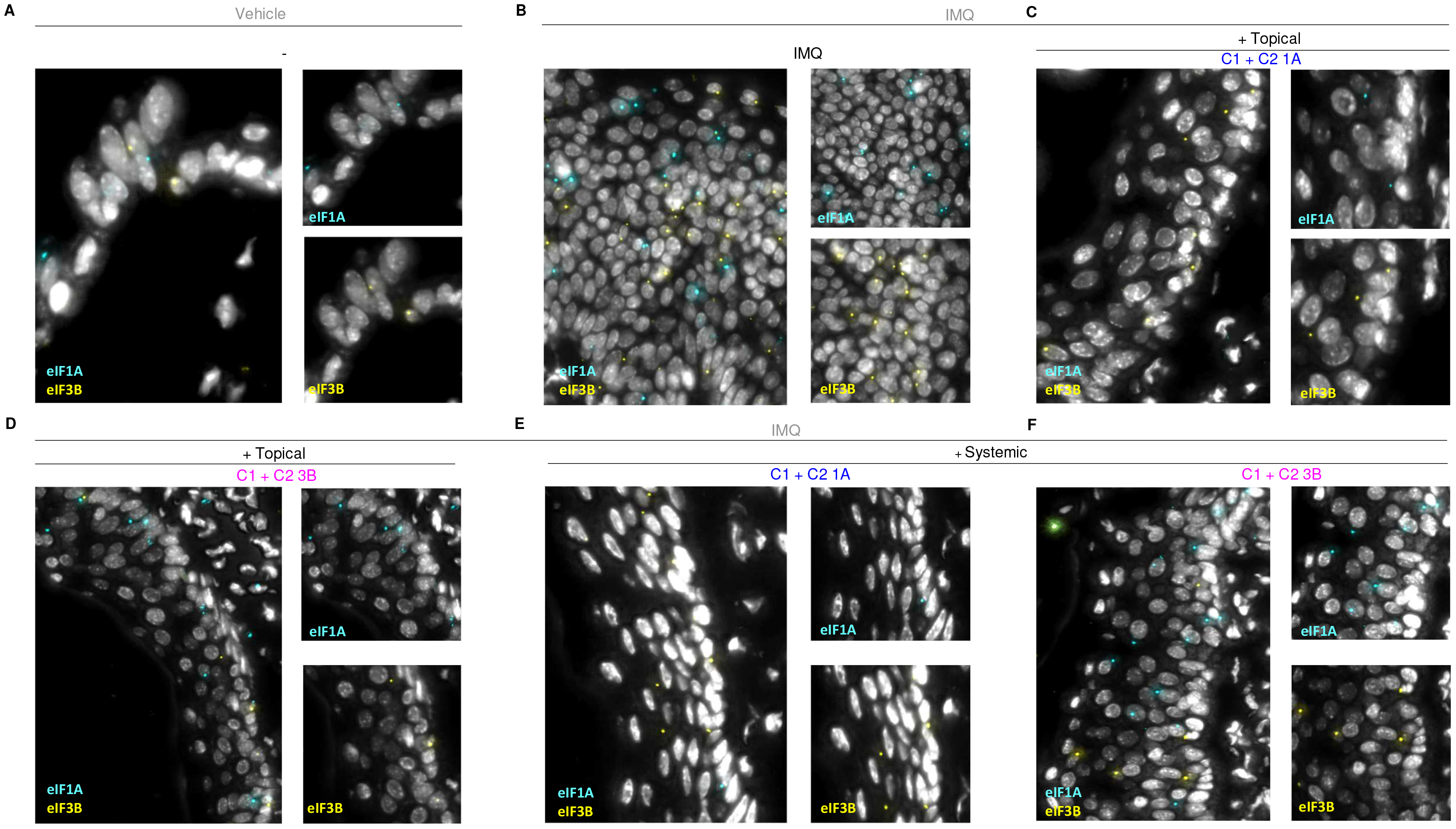
In situ mRNA detection of eIF1A and eIF3B by using padlock probes. (A - F) Representative images showing colocalization of eIF1A and eIF3B in keratinocytes in the IMQ mouse model. Nuclei are stained with DAPI and shown in white.

**Figure 5.**
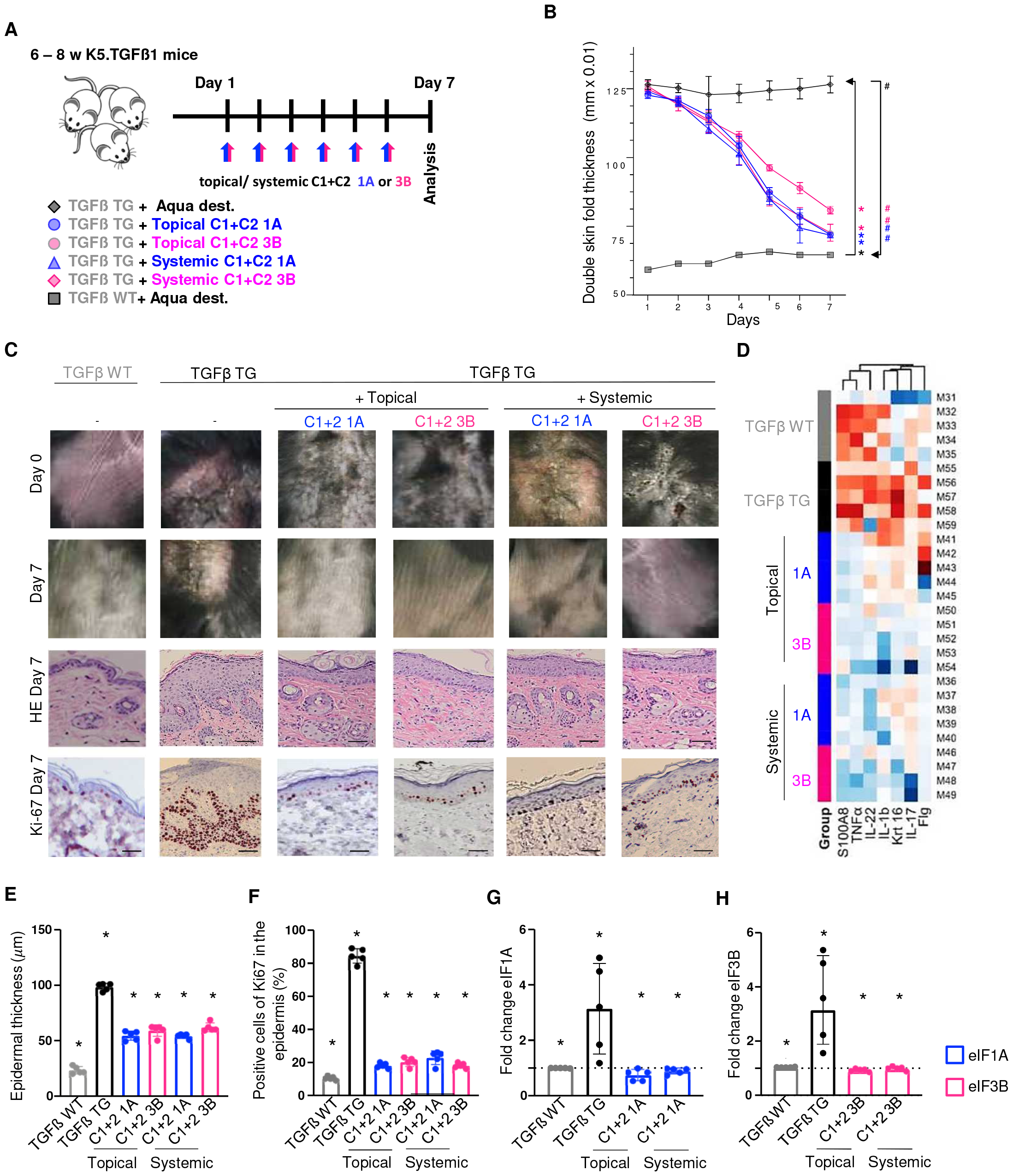
Topical and systemic siRNA treatment with eIF1A and eIF3B siRNA ameliorates the psoriatic phenotype in TGFß mice. **(A)** Topical and systemic administration of eIF1A and eIF3B siRNA for seven consecutive days. **(B)** Double skinfold thickness over the time of topical and systemic eIF1A and eIF3B siRNA application. **(C)** H & E and Ki-67 staining after topical or systemic eIF1A and eIF3B siRNA treatment on day 7. **(D)** Heat map of expression of proinflammatory cytokines and keratinocyte markers (ΔΔCT from RT-PCR, z-scaled per cytokine, normalized to reference group TGFß) shown with dendrograms sorted by average distance from dendrograms of hierarchical cluster analysis. **(E)** Epidermal thickness. **(F)** Ki-67 positive stained cells in the epidermis. **(G)** mRNA expression of *eIF1A*. **(H)** mRNA expression of *eIF3B. N* = 5 per group. ^*^ *p* < 0.05; ^**^ *p* < 0.01; ^***^ *p* < 0.001; ^****^ *p* < 0.0001.

**Figure 6.**
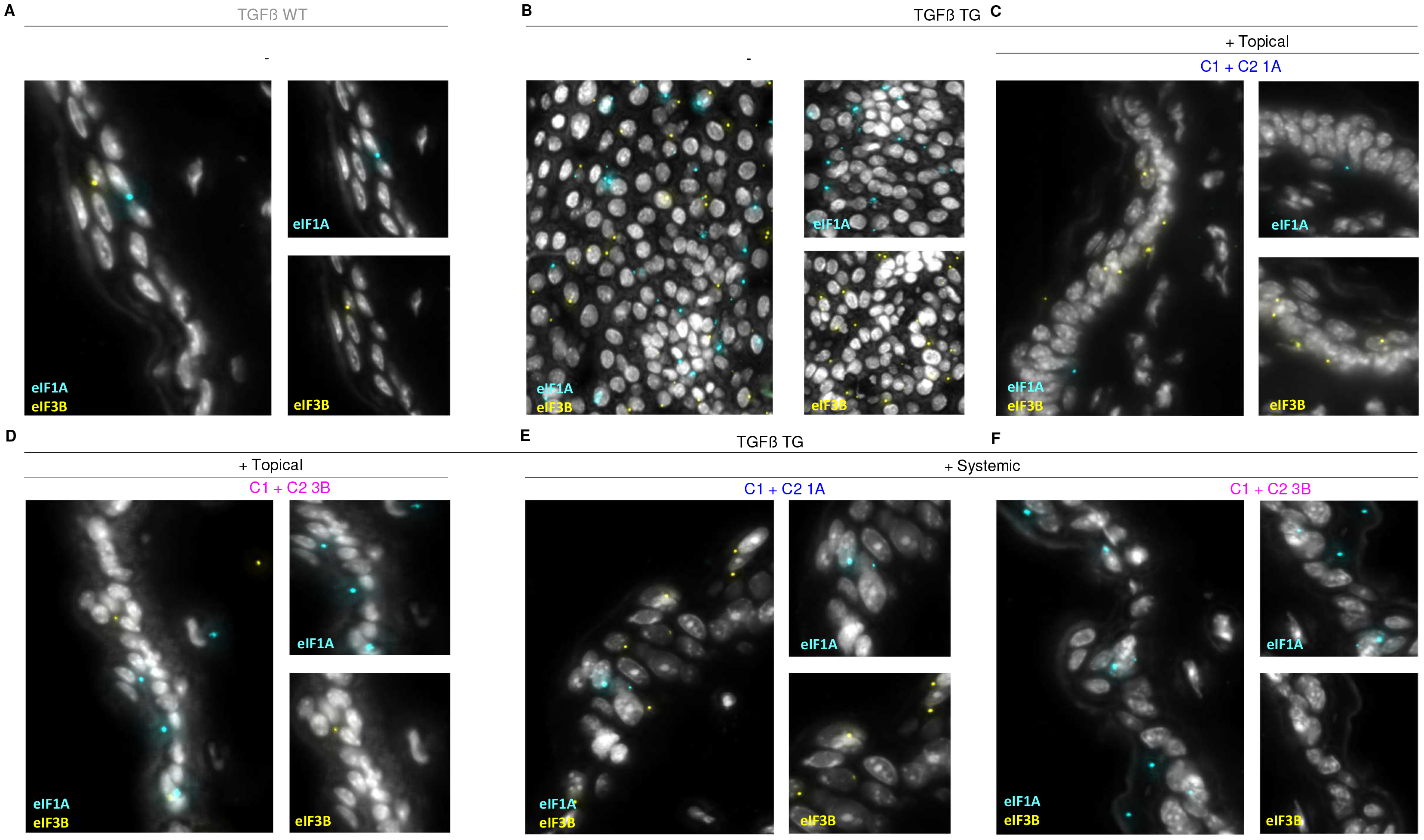
In situ mRNA detection of eIF1A and eIF3B by using padlock probes. (A - F) Representative images showing colocalization of eIF1A and eIF3B in keratinocytes in the TGFß mouse model. Nuclei are stained with DAPI and shown in white.

## Discussion

The regulation of gene expression in eukaryotes can occur at different stages, including mRNA translation. Eukaryotic mRNA translation is a complex process consisting of four main stages: Initiation, elongation, termination and ribosome recycling, with regulation occurring mainly in the initiation phase, which is the rate-limiting step of protein synthesis [20]. The most important regulators in the initiation phase are the eukaryotic translation initiation factors (eIFs). In eukaryotes, the 43S ribosomal preinitiation complex (PIC) consists of the 40S ribosomal subunit, the ternary eIF2-GTP-initiating methionyl-tRNA complex (Met-tRNAi) and many other eIFs such as eIF1, eIF1A, eIF3 and eIF5. The complex is recruited to the 5’-terminus of mRNAs and presumably scans the 5’-untranslated region (5’UTR). The PIC then moves to the start codon where the 60S ribosomal subunit joins the complex, leading to the formation of the 80S initiation complex. Subsequently, the 80S complex is able to recruit the correct aminoacyl-tRNA to the A (aminoacyl) site, which promotes the synthesis of the first peptide bond and shifts the initiation towards the elongation step [30].

The aim of this study was therefore to investigate the biological functions and pathophysiologic role of eIF1A and eIF3B in the development of psoriasis. Inhibition of eIF1A and eIF3B by siRNA resulted in abrogation of the psoriatic phenotype *in vivo* in IMQ and TGFß psoriasis models and *in vitro* cell culture studies. In addition, eIF1A and eIF3B by siRNA reduced the psoriatic phenotype in the human 3D model. Topical and systemic application of specific eIF1A and eIF3B siRNA significantly decreased the mRNA expression of eIF1A and eIF3B in the human 3D psoriasis model and in the IMQ and TGFß mouse models (Figure 2 - 4). It also normalized keratinocyte proliferation and keratinocyte differentiation molecules and disrupted the proinflammatory cytokine milieu by downregulating TNFα, IL-17 and IL-22. In addition, the application significantly improved the psoriasis phenotype at macroscopic and microscopic levels. Whether the effect of topical and systemic application of eIF1A or eIF3B on the inflammatory infiltrate in the dermis is direct or indirect remains to be determined.

Taken together, our work opens up several possibilities for potentially new topical and systemic treatment strategies for psoriasis. Not only by targeting the eIF signaling pathway, but also the AMPs and proinflammatory cytokines produced by keratinocytes are promising approaches. Our work suggests that topical and systemic siRNA silencing of eIF1A and eIF3B could be an efficient approach for the treatment of inflammatory skin diseases. Developing small molecule inhibitors of eIFs could be promising for the topical treatment of psoriasis.

## Material and Methods

### Animals

Mice (6-8 weeks old) were kept with water and food *ad libitum* in the conventional animal facility or a special pathogen-free animal facility at the Centre for Medical Research, Medical University of Graz, Austria. During the experiments, all mice were closely monitored to ensure their adequate health status. Female BALB/c mice were purchased from Charles River (Sulzfeld, Germany) and received a daily dose of 62.5 mg IMQ cream (5%) (MEDA Pharmaceuticals, Vienna, Austria) on the shaved back for 7 consecutive days. The TGFß model was generated by inserting 1.6 kb of full-length human TGFss1 wild-type cDNA into the K5 expression vector (63). Breeding pairs of K5.TGFβ1 transgenic mice were kindly provided by Prof. Xiao-Jing Wang (University of Colorado Anschutz medical campus, Colorado, USA) and bred at the Centre for Medical Research, Medical University of Graz, Austria. Twenty-four hours after the last experimental treatment, the mice were sacrificed, and samples were collected for further analysis. All animal experiments were performed in accordance with institutional guidelines and federal guidelines and were approved by the Austrian government, the Federal Ministry of Education, Science, and Research (protocol numbers: BMWF-66.010/0032-II /3b/2013 and 2021-0.393.968).

### Toxicity and safety of eIF1A and eIF3B siRNA

In all experiments, topical and systemic siRNA treatment was well tolerated and did not lead to local systemic changes. Overall, eIF1A and eIF3B siRNA appear to offer a treatment approach that is both effective and safe (Supplementary Figure 1).

### Histology

Paraffin-embedded human, mouse and 3D model samples were sectioned (3.5 µm) and stained with hematoxylin and eosin (H&E). The epidermal thickness was determined at five randomly selected sites per H&E sample under the microscope at 20x magnification. All measurements were performed in a blinded fashion. Finally, the results were averaged per human/mouse and per treatment group for statistical analysis. Staining images were acquired with an Olympus BX41 microscope (Olympus Life Science Solution, Hamburg, Germany) at 20x and 40x magnification using cellSens software (Olympus Life Science Solution, Hamburg, Germany).

### Immunohistochemistry

FFPE tissue sections (3.5 µm) were deparaffinized and rehydrated for immunohistochemical staining. Slides with tissue sections were incubated in Dako Target Retrieval Solution pH 9.0 (Dako S2367) for 30 minutes in a steam bath for heat-induced antigen retrieval. Staining was then performed manually at 4 °C by antibody incubation with the Dako REAL TM Detection System, Peroxidase/ACE, and antibodies against eIF1A (1:100; # ab177939; Abcam; Cambridge, UK) and eIF3B (1: 100; # ab124778, Abcam, Cambridge, UK) and Ki-67 (1:250; #12202S, Cell Signaling, Massachusetts, USA). As a secondary method, antibody staining was performed using a rabbit-specific HRP detection IHC kit (ab80436, Abcam, Cambridge) according to the manufacturer’s instructions. Ki-67- and eIF4E-positive cells were counted in a blinded manner in five randomly selected microscopic fields at 20x magnification.

### RNA extraction

Total RNA was isolated from materials frozen in RNAlater solution (Invitrogen, California, USA) at -80 °C from HaCaT cells, lesional skin from psoriasis patients, healthy skin from control subjects, human 3D psoriasis tissue culture and mouse skin. RNA was extracted using the phenol-chloroform extraction method. Cells and tissues were lysed in Trizol® reagent (Thermo Fischer Scientific Inc., Massachusetts, USA) before chloroform was added. After centrifugation, the aqueous phase was precipitated with isopropanol. The pellet was washed twice with 80 % alcohol and dissolved in RNase-free water. The RNA concentration in the supernatant was determined using a NanoDrop1000 spectrophotometer (Thermo Fischer Scientific, Massachusetts, USA).

### Quantitative real-time PCR

200 ng of RNA was reverse transcribed using a High Capacity cDNA Reverse Transcription Kit (Applied Biosystems, Foster City, USA) according to the manufacturer’s instructions. The cDNA was diluted 1:10 in RNase/DNAse-free water for quantitative real-time PCR (qRT-PCR). The qRT-PCR was performed using Power SYBR Green PCR Master Mix (Applied Biosystems, Foster City, USA) on a CFX96 Touch Real-Time PCR Detection System (Bio-Rad, Vienna, Austria). GAPDH was used as a stable housekeeping gene (HKG). The list of primer sequences is shown in Supplementary Table 1. Fold-change values were analyzed using the 2^-∆∆CT^ method.

### Cell culture

The human keratinocyte cell line HaCaT was obtained from the American Type Culture Collection (ATCC) and maintained in DMEM medium supplemented with 10% fetal bovine serum (FBS) and penicillin/streptomycin (100 µg/ml) and incubated in a humidified atmosphere with 5% CO2 at 37 °C. The cells were subjected to regulatory monitoring and confirmed to be free of mycoplasma.

### siRNA transfection

Transfection was performed using MetafecteneRsi+ transfection reagent (Biontex, Munich, Germany) according to the manufacturer’s instructions. The two human eIF4E small-interfering RNA (siRNA) sequences (eIF1A - C 1: TTGGATAAATACCTCGGACAT; eIF1A-C 2. AAGCCTGTAAACTATGCATGA; eIF3B – C 1: CGGGAAGATTGAACTCATCAA; and eIF3B – C 2: GACCGACTTGAGAAACTCAAA) were synthesized by QIAGEN (Hilden, Germany). For transfection, 1x SI buffer, Metafectene SI +, and 20 nM siRNA (eIF4E-1, eIF4E-2, and both eIF4E siRNAs together) were mixed to one drop per well. After incubation for 15 min at room temperature, 2 x 10^4^ cells per well were seeded onto a 6-well plate. The cells with the transfection mixture were cultured at 37°C in a humidified atmosphere with 5% CO2. The cells were harvested 24 h, 48 h, and 72 h after transfection.

### Proliferation assay

Transfected cells were seeded in 96-well plates (8 x 104 cells/well) and cultured for 24 h, 48 h and 72 h. The number of viable cells was determined. The number of viable cells was determined by mitochondrial conversion of 3-(4, 5-dimethylthiazol-2-yl)-2,5-diphenyltetrazolium bromide (MTT) (Sigma Aldrich, Missouri, USA) to formazan. The cells were incubated with MTT for 2 hours at 37 °C, the supernatant was removed, and the cells were lysed with sodium dodecyl sulfate and isopropanol/HCl with shaking for 15 minutes at room temperature. The optical density was measured at 570 nm (SynergyTM4, BioTek, Winooski, USA).

### Human 3D psoriasis tissue model

Human 3D psoriasis tissue models were purchased from MatTec (MatTek Corporation, Massachusetts, USA). The model was treated according to the manufacturer’s instructions. The psoriatic tissue models were treated by epicutaneous application of or by adding equal amounts to the cell culture medium. One part was fixed in 4% paraformaldehyde, the other part was frozen in liquid nitrogen and stored until the RNA and protein were extracted.

### In situ sequencing

In situ sequencing was performed using an in situ sequencing (ISS) method based on cDNA hybridization (pubmedID: 32990747). The ISS procedure included three cycles for specific detection and visualization of Eif1ax and Eif3b as well as transcripts. Padlock probes were designed using the open-source Python-based padlock design software, which was packaged as previously described (pubmedID: 32990747). We used the following GenBank accession numbers NM_025437.5 for Eif1ax and NM_133916.2 for Eif3b, which we obtained from the National Center for Biotechnology Information (NCBI). In brief, cDNA synthesis was initiated with target-specific reverse transcription primers. Padlock probes were then hybridized to the cDNA, followed by ligation. The circularized padlock probes were then amplified by rolling circle amplification. The Eif1ax and Eif3b transcripts were visualized using Cy7 as an anchor and detected in specific channels. All sequences are listed in the attached Supplementary Table 2.

### General statistical analyses

Statistical analyzes were performed using GraphPad Prism version 8 (GraphPad Software, California, USA). Data were tested for normal distribution and similar variances between groups. Statistical significance was assessed using the two-tailed, unpaired Student’s test for comparisons between two groups or one-way ANOVA with appropriate post hoc tests for multiple comparisons. If the data were not normally distributed or showed unequal variance between two groups, the two-tailed Mann-Whitney U test was used for statistical analysis. Statistical significance was set at *p* ≤ 0.05. Heat maps of RT-PCR expressions (∆∆CT z-scaled per cytokine, normalized to reference group) were created in R v4.2.1 with *pheatmap::pheatmap()*. Cytokines were clustered by Lance-Williams dissimilarity update with complete linkage using *stats::dist()* and *stats::hclust()* and dendograms were sorted with *dendsort::dendsort()* at every merging point according to the average distance subtrees.

## Author contributions

Conceptualization, NGS, PW; Methodology, NGS, JP, AV, NS, CTM, AEH; Validation, NGS, PW, NS; Formal analysis, NGS, NB, CTM, AEH; Investigation, NGS; Resources, all authors; Data curation, NGS, NB, MCT; Writing – original draft, NGS, PW; Writing – Review & Editing, all authors; Visualization, NGS, NB, CTM, AEH; Supervision, NGS, PW; Project administration, NGS; Funding acquisition, NGS, PW.

## Acknowledgements

The study was supported by Austrian Science Fund FWF W1241 to PW and the PhD program Fundamentals in Molecular Inflammation (MOLIN), and the Cultural Office of the City of Graz and the Scientific Association of Styrian Dermatology. The authors are grateful to Sara Crockett for editing the manuscript and Peter Nussbaumer, Andreas Billich, and Wolfgang Sommergruber (Wings of Innovation, Austria) for support and discussion.

