## Supplemental Table 1 for "Inhibition of eukaryotic translation initiation factor 1A (eIF1A) and 3B (eIF3B) diminishes the psoriatic phenotype in two mouse models and human 3D model samples"

Golob-Schwarzl et al., Table S2
Primer sequences for qRT-PCR

| **Gene** | **Primer Pair** | **Sequence (5’-3’)** | **Tm [°C]** |
| --- | --- | --- | --- |
| **eIF1A** | Fwd | GAAACGGCAGGAAGACCCTTA | 60 |
|  | Rev | CGGATGCTCAATTACAGTACCAT | 59 |
| **eIF3B** | Fwd | GGACCCGACCGACTTGAGA | 63 |
|  | Rev | TTGACCCGGAATGTGTGCTG | 63 |
|  | Rev | CGCCCTCGAACACACTGTAGAAGT | 64 |
| **Il-17** | Fwd | GGACTCTCCA CCGCAATGA | 60 |
|  | Rev | TCAGGCTCCCTCTTCAGGAC | 60 |
| **IL-22** | Fwd | CAGCTCCTGTCACATCAGCGGT | 63 |
|  | Rev | AGGTCCAGTTCCCCAATCGCCT | 63 |
| **S100A8** | Fwd | AAATCACCATGCCCTCTACAAG | 58 |
|  | Rev | CCCACTTTTATCACCATCGCAA | 58 |
| **Flg** | Fwd | GAAGGAACTTCTGGAAGGACAAC | 60 |
|  | Rev | TCCATCAGTTCCACCATGCCTC | 60 |
| **Krt** | Fwd | AGCAGGAGATCGCCACCTA | 60 |
|  | Rev | AGTGCTGTGAGGAGGAGTGG | 60 |
| **IL-1b** | Fwd | GAGTGTGGATCCCAAGCAAT | 58 |
|  | Rev | TACCAGTTGGGGAACTCTGC | 58 |
