## Supplemental Table 2 for "Inhibition of eukaryotic translation initiation factor 1A (eIF1A) and 3B (eIF3B) diminishes the psoriatic phenotype in two mouse models and human 3D model samples"

| **Primers** | **Sequences (5´ - 3´)** |
| --- | --- |
| RV_Eif1ax_1 | CAGGAACTGCGGTGA |
| RV_Eif1ax_2 | GCAAAGACTGGCTGG |
| RV_Eif1ax_3 | CGCTTTTCAGATTCA |
| RV_Eif1ax_4 | ATCCTCCTTGAACACA |
| RV_Eif1ax_5 | AAAATGACATCGGCT |
| RV_Eif1ax_6 | CATCTGCGTTGTATT |
| RV_Eif1ax_7 | GGACCAAATGTATCAG |
| RV_Eif1ax_8 | CCCAATGTCATCAAACT |
| RV_Eif1ax_9 | AGGTATGGTGGTACA |
| RV_Eif1ax_10 | GACCTATCATTTTTGCC |
| RV_Eif1ax_11 | CTCCAAACCCAAAAAACCAT |
| RV_Eif1ax_12 | CCGCCCACAAACAAA |
| RV_Eif1ax_13 | GCATGAAAAACCAGT |
| RV_Eif1ax_14 | GCAACTACTGACCTTA |
| RV_Eif1ax_15 | CAGACGACAAGATCC |
| RV_Eif1ax_16 | CCCCACGGTAAACTT |
| RV_Eif1ax_17 | GATGGATCTTGGTTTGA |
| RV_Eif1ax_18 | AATACAAGGTCCGCA |
| RV_Eif1ax_19 | CAGATCTCCTTGTTTGT |
| RV_Eif1ax_20 | GGAGATGAAGTCATAGT |
| RV_Eif3b_1 | TTCGGCGTCTTCCGT |
| RV_Eif3b_2 | TTGGCCCTCACTTCG |
| RV_Eif3b_3 | GCTCATCTGCCTCTC |
| RV_Eif3b_4 | CGAAATCCTCGGGGT |
| RV_Eif3b_5 | CTTTGAGTACATCACC |
| RV_Eif3b_6 | AATCCCGTCCGCTTC |
| RV_Eif3b_7 | CAGCATCCACAGCAT |
| RV_Eif3b_8 | CATCGGCGTTTTTCA |
| RV_Eif3b_9 | ACACTGTACTGATCTCT |
| RV_Eif3b_10 | CCGCTCTCAAAAATC |
| RV_Eif3b_11 | GCTGATGGAAAGTGG |
| RV_Eif3b_12 | AACTTGTCCCCTCCC |
| RV_Eif3b_13 | AGTGGGCTGAGCTTT |
| RV_Eif3b_14 | ATGGCTCCACTTGAA |
| RV_Eif3b_15 | CCTCACTCGGATCTCCT |
| RV_Eif3b_16 | CAACCACATTGAACA |
| RV_Eif3b_17 | ACGTGGTAGAAGGAGA |
| RV_Eif3b_18 | TTCCCGTTGCTCTTC |
| RV_Eif3b_19 | CAATGTTCATGACGG |
| RV_Eif3b_20 | GGAGGCCATGTAGTG |
| **Padlock probes** | **Sequences (5´ - 3´)** |
| plp_Eif1ax_1 | /5Phos/TTCTCGGCGTTGCCC**TCTACGAGTTTGCAGTCACGTGCGTCTATTTAGTGGAGCC**TACTTGTCACTCAGACCGTCTTCGTGGCGCGCAC |
| plp_Eif1ax_2 | /5Phos/TAGAGGCAAAAATGA**TCTACGAGTTTGCAGTCACGTGCGTCTATTTAGTGGAGCC**TACTTGTCACTCAGACCGTAGGCAAAAACCGGCG |
| plp_Eif1ax_3 | /5Phos/AGGACAACAAAGCCG**TCTACGAGTTTGCAGTCACGTGCGTCTATTTAGTGGAGCC**TACTTGTCACTCAGACCGTGGCTACGAGATTACC |
| plp_Eif1ax_4 | /5Phos/CATGCTAAAATCAAT**TCTACGAGTTTGCAGTCACGTGCGTCTATTTAGTGGAGCC**TACTTGTCACTCAGACCGTGGCGAGCTTCCAGAG |
| plp_Eif1ax_5 | /5Phos/TGAGTGCTAAGGGCA**TCTACGAGTTTGCAGTCACGTGCGTCTATTTAGTGGAGCC**TACTTGTCACTCAGACCGTTTGCTGGGATCCTCC |
| plp_Eif1ax_6 | /5Phos/TGGAGGAGGAAAGGC**TCTACGAGTTTGCAGTCACGTGCGTCTATTTAGTGGAGCC**TACTTGTCACTCAGACCGTCAGTCATCGGCTTAC |
| plp_Eif1ax_7 | /5Phos/AAAATAGTTTTCGAA**TCTACGAGTTTGCAGTCACGTGCGTCTATTTAGTGGAGCC**TACTTGTCACTCAGACCGTGCACGGCCCGTCTCC |
| plp_Eif1ax_8 | /5Phos/GTAGTGTGTCGCAGA**TCTACGAGTTTGCAGTCACGTGCGTCTATTTAGTGGAGCC**TACTTGTCACTCAGACCGTGCTATAGCCGAAAAA |
| plp_Eif1ax_9 | /5Phos/ACTAGACATGCTGAA**TCTACGAGTTTGCAGTCACGTGCGTCTATTTAGTGGAGCC**TACTTGTCACTCAGACCGTAGCTAGTCGGAACTC |
| plp_Eif1ax_10 | /5Phos/ACGTGGTTTCACAAA**TCTACGAGTTTGCAGTCACGTGCGTCTATTTAGTGGAGCC**TACTTGTCACTCAGACCGTAGGAATGCGCCTTTC |
| plp_Eif3b_1 | /5Phos/GCCGACGGACGAGGC**CCTCAATGCTGCTGCTGTACTACTGCGTCTATTTAGTGGAGCC**TCGTATAGCTGTATCGGGCCTCCGAGTCGCC |
| plp_Eif3b_2 | /5Phos/CTGGAGCGGAGGAGG**CCTCAATGCTGCTGCTGTACTACTGCGTCTATTTAGTGGAGCC**TCGTATAGCTGTATCGGGGGGCACCCCTCGG |
| plp_Eif3b_3 | /5Phos/GGAAGAACTACTTGG**CCTCAATGCTGCTGCTGTACTACTGCGTCTATTTAGTGGAGCC**TCGTATAGCTGTATCGAGACGACGTGAGCGA |
| plp_Eif3b_4 | /5Phos/GTACGCTTCTCCCGC**CCTCAATGCTGCTGCTGTACTACTGCGTCTATTTAGTGGAGCC**TCGTATAGCTGTATCGGTACATCTTTCTGGA |
| plp_Eif3b_5 | /5Phos/GGAAGAGGCAGAGTG**CCTCAATGCTGCTGCTGTACTACTGCGTCTATTTAGTGGAGCC**TCGTATAGCTGTATCGTTTGCGTTACTGGCT |
| plp_Eif3b_6 | /5Phos/CCCCTAAGGGTACCT**CCTCAATGCTGCTGCTGTACTACTGCGTCTATTTAGTGGAGCC**TCGTATAGCTGTATCGCGTACGTGCGCTGGT |
| plp_Eif3b_7 | /5Phos/CATAAGAAGAGAGGT**CCTCAATGCTGCTGCTGTACTACTGCGTCTATTTAGTGGAGCC**TCGTATAGCTGTATCGGACATCCTCACGGGC |
| plp_Eif3b_8 | /5Phos/ATGCAGCTCCCCACC**CCTCAATGCTGCTGCTGTACTACTGCGTCTATTTAGTGGAGCC**TCGTATAGCTGTATCGGCCAGGGTGACCCTG |
| plp_Eif3b_9 | /5Phos/CCCTCGGATCTCTGT**CCTCAATGCTGCTGCTGTACTACTGCGTCTATTTAGTGGAGCC**TCGTATAGCTGTATCGGCTTCATGGTGAGGC |
| plp_Eif3b_10 | /5Phos/TTCAGACTGCACCGT**CCTCAATGCTGCTGCTGTACTACTGCGTCTATTTAGTGGAGCC**TCGTATAGCTGTATCGGGCGTTTGTCGACAC |
| **Detection probes** | **Sequences (5´ - 3´)** |
| D1 | TexasRed-TCTACGAGTTTGCAGTCACG |
| D2 | Cy3-CCTCAATGCTGCTGCTGTACTAC |
| D3 | Cy3-TCTACGAGTTTGCAGTCACG |
| D4 | CY5 CCTCAATGCTGCTGCTGTACTAC |
| D5 | CY7-UGCGUCUAUUUAGUGGAGCC |
| Oligonucleotide sequences. Padlock probes were 5’-phosphorylated. |  |
| The detection probes were 5´conjugated with fluorophores (Texas Red, CY3, CY5 and CY7 are fluorescent labels). |  |
| +: underlined: target complement sequence, bold: detection probe complement sequence. |  |
