## Supplementary figures and images for "Inhibition of eukaryotic translation initiation factor 1A (eIF1A) and 3B (eIF3B) diminishes the psoriatic phenotype in two mouse models and human 3D model samples"

### Supplemental Figure 1

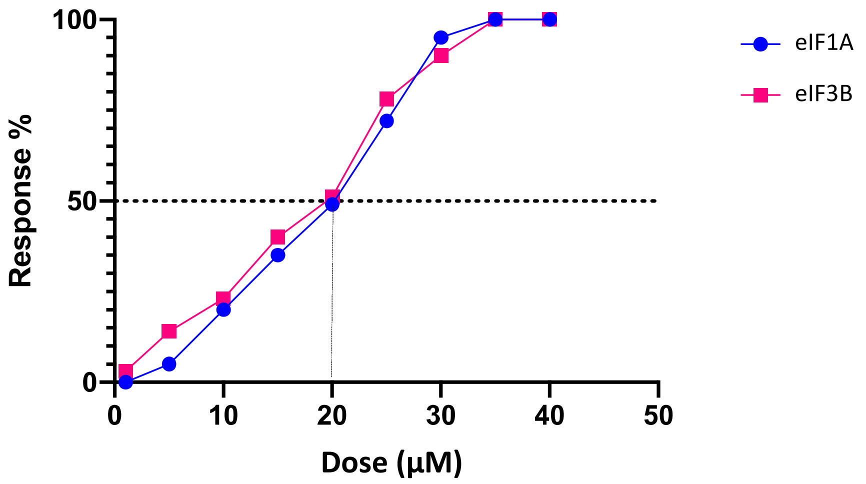
